# Comparative Multi-scale Hierarchical Structure of the Tail, Plantaris, and Achilles Tendons in the Rat

**DOI:** 10.1101/396309

**Authors:** Andrea H. Lee, Dawn M. Elliott

## Abstract

Rodent tendons are widely used to study human pathology, such as tendinopathy and repair, and to address fundamental physiological questions about development, growth, and remodeling. However, how the gross morphology and the multi-scale hierarchical structure of rat tendons, such as the tail, plantaris, and Achillles tendons, compare to that of human tendons are unknown. In addition, there remains disagreement about terminology and definitions. Specifically, the definition of fascicle and fiber are often dependent on the diameter size and not their characteristic features, which impairs the ability to compare across species where the size of the fiber and fascicle might change with animal size and tendon function. Thus, the objective of the study was to select a single species that is widely used for tendon research (rat) and tendons with varying mechanical functions (tail, plantaris, Achilles) to evaluate the hierarchical structure at multiple length scales. This study was designed including, histology, SEM, and confocal imaging. We confirmed that rat tendons do not contain fascicles, and thus the fiber is the largest tendon subunit in the rat. In addition, we provided a structurally-based definition of a fiber as a bundle of collagen fibrils that is surrounded by elongated cells, and this definition was supported by both histologically processed and unprocessed tendons. In all rat tendons studied, the fiber diameters were consistently 10-50 µm, and this diameter appears to be conserved across larger species. Specific recommendations were made for the strengths and limitations of each rat tendon as tendon research models. Understanding the hierarchical structure of tendon can advance the design and interpretation of experiments and development of tissue engineered constructs.

## Introduction

Rodent tendons are widely used to study human pathology, such as tendinopathy and repair, and to address fundamental physiological questions about development, growth, and remodeling. The rat tail tendon is widely used because its structure is relatively simple; however, rat tail tendon bears low stresses and lacks muscle-tendon junctions.^1,2^ The rat plantaris and Achilles tendons are also popular model systems^3–8^ and there is an increasing interest in the plantaris tendon as it has recently been suggested to contribute to chronic Achilles tendinopathy.^9–11^ However, the gross morphology and the multiscale hierarchical structure of these rat tendons are not fully understood.

Tendon hierarchical structure, spanning multiple length scales, is widely known, yet there remains disagreement about terminology and definitions, which hinders communication among scientists and interpretation of experimental results. For example, much of the literature uses the terms fiber and fiber bundle to generally refer structures made of collagen. This general usage of “fiber” can cause confusion when studies report fibrils^12^ and fascicles^13^ as fibers. It is widely accepted that the fascicle is the largest tendon subunit,^13–15^ followed by smaller subunits called fibers, and each fiber is composed of fibrils. However, the definition of fascicle and fiber are often dependent on the diameter size and not their characteristic features, which impairs the ability to compare across species where the size of the fiber and fascicle might change with animal size. In large animals, the surrounding interfascicular matrix boundaries are easily visible^16^, yet a similar structure cannot be found in rat tail tendon, despite the fact that the term ‘fascicle’ is routinely used to describe a rat tail tendon. The fascicle should be defined as a bundle of fibers with distinct interfascicular matrix boundaries that is populated with round cells.^17,18^ Because the fascicle size in larger animals^19^ is about the same size of an entire tendon in a rat, it is unclear if rat tendon has fascicles. We hypothesize that rat tendons do not have fascicles due to its small size. Precise definition and terminology of these hierarchical structures are crucial because the structures are directly related to tendon physiology and mechanical function.

Another source of confusion is the definition, or even existence, of the fiber. While some include the fiber as a part of tendon hierarchical structure schematics^14,20,21^, others exclude it,^13,22^ and the terminology of fiber is used inconsistently. Moreover, when fibers are defined, it is done by diameter size, which may be species-dependent.^19,23^ This definition is problematic, and it is hard to distinguish between a fiber from fascicle because they have a large and overlapping size ranges. ^20,21^ An alternative way to define a fiber is based on another structural component such as cells. Elongated tendon cells reside parallel to the direction of fibrils and a group of cells is aligned and connected by gap junction.^24–26^ These cells outline a bundle of collagen fibrils, and this bundle may be a unit of fiber. This definition is consistent with the developmental process of tendon, where multiple cells in contact deposit collagen between cell-cell junctions during early development.^27^ This process leads to cells surrounding a bundle of collagen fibrils at later stages of development^27^, and thus this supports the hypothesis that cells may define the boundary of a fiber. In this study, we will test our hypothesis that fiber is defined by surrounding cells.

Tendon hierarchical structures may also depend on loading environment. For example, there are differences between low- and high-stress bearing tendons for multi-scale mechanics^16,28–30^ and composition^31^ in equine and bovine tendons. These differences between tendons with different mechanical functions suggest that their hierarchical organizations may be different. ^19^ Thus, we selected the tail, plantaris, and Achilles tendons to represent tendons with different mechanical functions.

The objective of this study was to evaluate the hierarchical structure of rat tendons at multiple length scales. Because the hierarchical structure is likely to depend on the species and loading environment, we chose a single species that is widely used for tendon research (rat) and tendons with varying mechanical functions (tail, plantaris, Achilles). This study was designed to investigate the gross morphology at the macro-scale and the hierarchical fascicle and fiber structures at the micro-scale by using multiple imaging methods including, histology, SEM, and confocal imaging. In addition, we reconstructed a three-dimensional (3D) structure from confocal imaging stacks. Understanding and properly defining tendon hierarchical structures are critical in research to advance the design and interpretation of experiments and development of tissue engineered constructs.

## Methods

### 1. Sample Preparation

All tendons were from 4-7-month-old female Long Evans rats and sacrificed on the day of the experiment. Rat tail (n=13), plantaris (n=13), and Achilles (n=13) tendons were divided into five experimental groups, where each group included different imaging technique. Each tendon was processed for longitudinal PicroSirus Red staining (n=3/tendon type), transverse H&E staining (n=3/tendon type), longitudinal scanning electron microscope (SEM) (n=3/tendon type), confocal imaging with second harmonic generation (SHG) signals and cells (n=3/tendon type), or 3D rendering of confocal image stack (n=1/tendon type).

### 2 Gross Morphology

Rat tail tendon was dissected following the same protocol from a published study.^32,33^ Plantaris and Achilles tendons were dissected using bone cutting shears, scalpels, and forceps to dissect the Achilles and plantaris complex about 4 cm in length with the calcaneus attached. We separated out both plantaris and Achilles from each other by removing excess surrounding soft tissue. Each tendon was randomly assigned to different imaging groups after the dissection. An additional Achilles tendon was further dissected using a sharp dissection microscope to study how Achilles tendon is fused. We separated each sub-tendon to verify that Achilles tendon is fused tightly. This sample was discarded after taking a reference picture (Fig 1) and was not used further in the study.

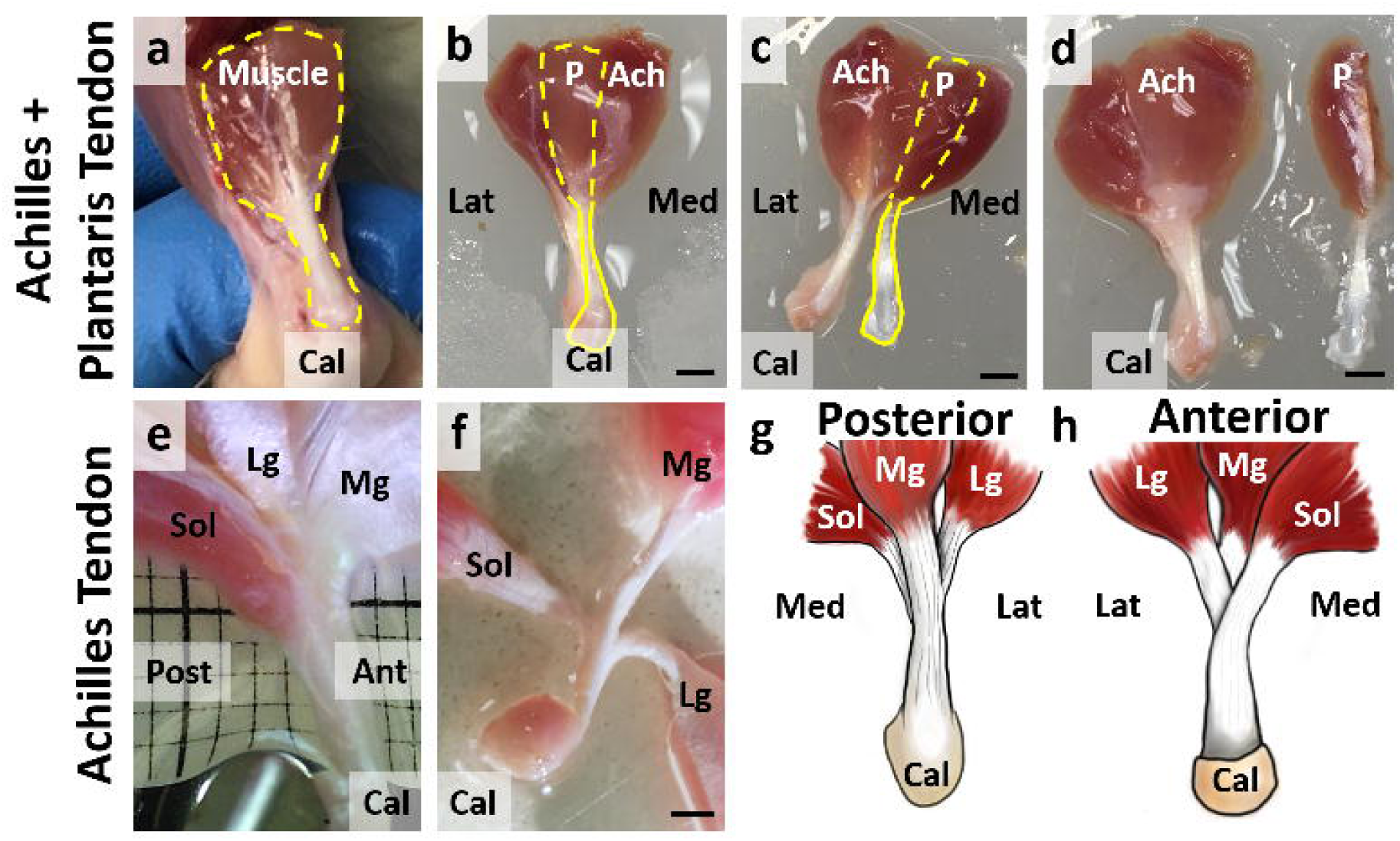

### 3 Histology

Histological analyses were performed in longitudinal and transverse directions to observe the hierarchical structure of tendons in both directions. Each tendon was fixed in 4% paraformaldehyde at room temperature. For the longitudinal histology, all tendons were fixed overnight. For transverse histology, the tail tendon was fixed for 30 minutes, the plantaris tendon was fixed for 2 hours, and the Achilles tendon was fixed overnight. The fixed tendons were then incubated for 30 minutes in a 30% sucrose solution to avoid possible ice damage. To process as frozen sections, each tendon was mounted on a plastic mold with optimal cutting temperature compound (OCT) and was flash frozen with an indirect contact with liquid nitrogen. We used CryoJane Tape Transfer System (Leica Biosystems) to cut 14 µm thickness sections for the longitudinal sections and 12 µm thickness sections for the transverse sections. We cut the mid-plane of the tendon, away from both anterior and superior planes. The longitudinal sections were stained with PicroSirus Red to visualize collagen alignment, and transverse sections were stained with H&E to visualize general organization. The staining process followed a standard staining procedure for frozen sections, and the slides were mounted to be visualized under Axio Imager 2 Pol (Carl Zeiss Inc). Images were acquired at two different magnifications (20x and 40x).

### 4 Scanning Electron Microscopy (SEM)

We imaged tendon structure using SEM to visualize the ultrastructure and confirm histological observations. Tendons used for SEM were processed following the same procedure as histology sections until staining (n=3/tendon). Using the cryostat, each tendon was cut once in a longitudinal direction of mid-plane to image the ultrastructure of mid-plane, and the sample was briefly placed in water to remove the remaining OCT. The section was processed for SEM using a standard processing technique consisting of fixation with 1% osmium tetroxide followed by a series of dehydration with ethanol. Each sample was dried using a critical point dryer, mounted on an SEM sample holder using carbon tabs, and coated with platinum (Leica Biosystems). The samples were visualized with a field emission SEM (Hitachi S-4700).

### 5 Confocal Imaging with Second Harmonic Generation (SHG) Signals and Cells

To test our hypothesis that cells outline fiber in tendon, we acquired SHG signals for collagen alignment and fluorescence from cell for cell placement. SHG microscopy is a nonlinear optical method that can visualize collagen without additional staining or processing.^34–37^ Immediately after the sacrifice and dissection, tendon was incubated in DMEM culture medium (Dulbecco’s Modified Eagle Medium) supplemented with 25 mM HEPES, 1% penicillin, and 1% streptomycin (ThermoFisher Scientific) for 30 minutes at 37C. Each tendon was stained with 10 µM CellTracker Green CMFDA dye (ThermoFisher Scientific) for 45 minutes and washed in the medium for 30 min at 37C. Each sample was placed in a petri dish to image with a laser-scanning multi-photon microscope (LSM 780, objective Plan-Apochromat 20x/1.0, Carl Zeiss Inc). We used Chameleon Vision II Ti:S laser (Coherent Inc.) tuned to 810 nm and an external detector to collect SHG signals. A cover glass was placed on top of the tendon to prevent it from floating. The confocal image stacks (0.42 × 0.42 × 0.67 μm pixel^-1^) were taken at the mid-substance of each tendon. The SHG signals of the most superficial (closest to the skin) and deep layers (furthest from the skin) were separated from the rest of the stack to show the change of collagen alignment through the depth. To show the cell placement, an optical slice that showed the strongest cell staining in the deep plane was separated from the confocal stack.

### 6 3D Rendering of Confocal Images

To observe collagen alignment and cell placement in 3D, a separate confocal image stack from above, spanning ∼50 μm in the z-direction, was acquired in the same way as the previous section in methods. The confocal image stacks were rendered by software AMIRA 6.3 (ThermoFisher Scientific). The blood vessels and cells were selected using an automatic threshold method under segmentation editor. Additional structures that were not captured by the automatic segmentation were selected manually. The surface of vessels and cells were rendered, and the volume of the SHG signal of collagen was rendered to show the 3D structure of tendon. The opacity of collagen was adjusted to show the 3D structure through the depth. A supplementary video of the 3D structure was created using AMIRA 6.3 animation function.

## Results

### Gross Morphology of Plantaris and Achilles Tendons

The gross morphology of plantaris (P) and Achilles (Ach) tendon demonstrated its structural complexities. Plantaris and Achilles tendons are bound together in vivo (Fig 1a). The plantaris muscle originates on the anterior-lateral side of the Achilles muscle, passes over the calcaneus (Cal), and inserts into the bottom of the foot. Once the calcaneus is cut, the anatomy of plantaris and Achilles tendons can be easily distinguished from each other. The plantaris muscle is behind Achilles muscle complex (Fig 1b, dashed yellow line), and the plantaris tendon is visible above Achilles tendon (solid yellow line). We separated the two tendons by cutting connective tissue binding the plantaris and Achilles tendons together at the insertion (Fig 1c, d). Human plantaris tendon has been previously described as a vestigial tissue that is absent for a large portion of the population.^38–41^ However, recent studies identified plantaris tendon in all or most of cadavers with variable insertion sites.^9,42,43^ We observed a single insertion sitefor the rat plantaris tendon, unlike human plantaris tendon. The Achilles tendon was further dissected to observe its gross morphology and verified that the Achilles tendon is a fusion of three tendons originating from soleus (Sol), lateral gastrocnemius (Lg), and medial gastrocnemius (Mg) (Fig 1e) muscles, similar to human.^44,45^ The sub-tendons from these muscles are tightly fused approximately at the half of the tendon length and cannot be further separated without directly cutting the tendon (Fig 1f). Among the three muscle groups, the Sol tendon is the most anterior sub-tendon, and Mg tendon is the most posterior sub-tendon. Schematics of the posterior (Fig 1g) and anterior (Fig 1h) views show the complex 3D morphological structure of Achilles tendon.

### Histological Evaluation of Tendon Micro-scale structure

To investigate the hierarchical structure and cell morphology in rat tendons, we conducted several analyses using multiple imaging methods. In longitudinal sections stained with PicroSirus Red, we observed a separation between structures with a diameter of 10-50 µm, at both low (Fig 2a-c) and high magnifications (Fig 2d-f) for all three tendons. The hierarchical structure defined by this unit of separation, highlighted in white arrows, is likely to be a fiber. Supporting this notion, we also observed the structures with a similar diameter in longitudinal view of SEM in all of the tendons, highlighted in green (Fig 2g-i). In addition, the transverse sections stained with H&E also had separations with a similar diameter at low (Fig 3a-c) and high magnifications (Fig 3d-f), outlined in blue. Rat tail tendon did not have a similar pattern of separation between the subunits as seen in plantaris and Achilles tendons in the transverse direction, perhaps due to its small overall size. Nonetheless, the distance between cells of tail tendon (blue arrows, Fig 3d) was similar to the separation observed in plantaris and Achilles tendons (blue outlines, Fig 3e,f).

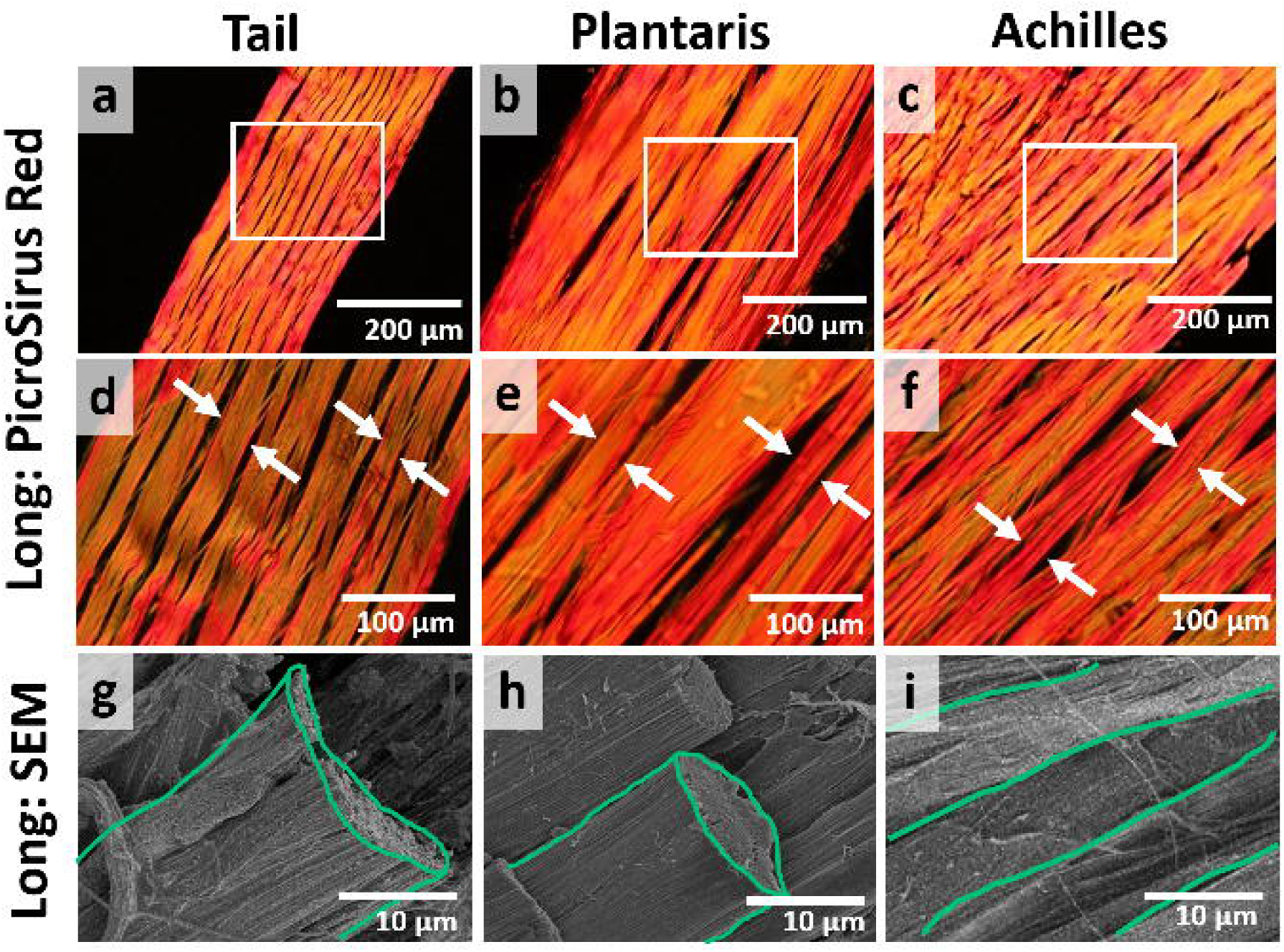

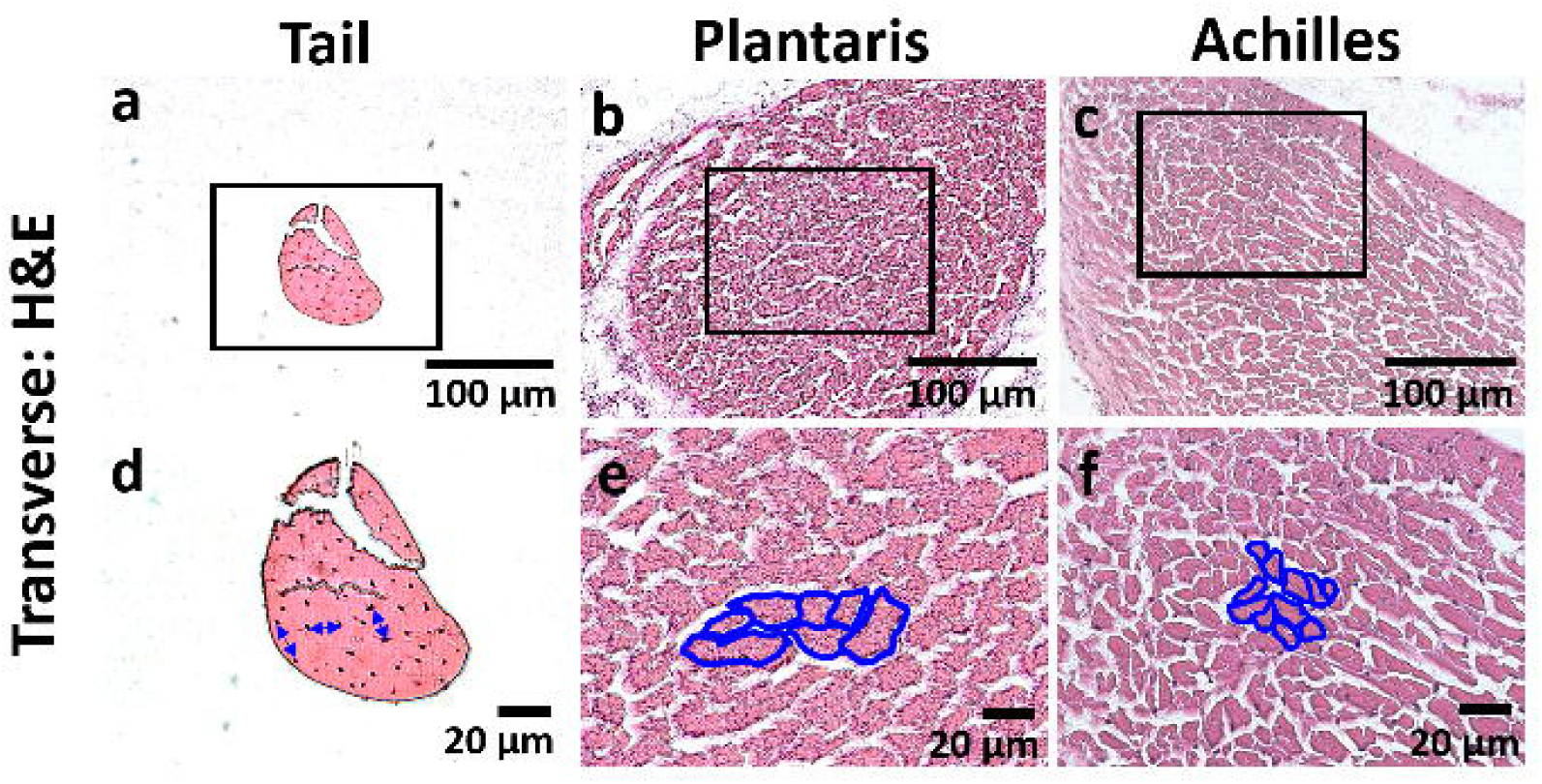

### SHG Evaluation of Fiber Structure and Cell Morphology

The collagen structure and cell morphology were visualized using the confocal microscope to confirm histological findings without the potential fixation and histological artifacts. At the deep plane (away from the surface of the tendon) all tendons had aligned collagen (Fig 4a-c). At the superficial plane (the surface of tendon) tail tendon had no change in collagen alignment from deep planes (Fig 4d), while both plantaris and Achilles tendons had an additional layer of peritenon, a complex meshwork of collagen that was about 20 µm thick (Fig 4e, f). In deep planes, the cells with elongated morphology aligned along collagen bundles (white arrows) that were the same diameter as observed histology (Fig 2, 3). Thus, rat tendon has a single hierarchical structure with a diameter of 10-50 µm that is surrounded by the elongated cell, and we define them as fiber from here and on (Fig 4g-i).

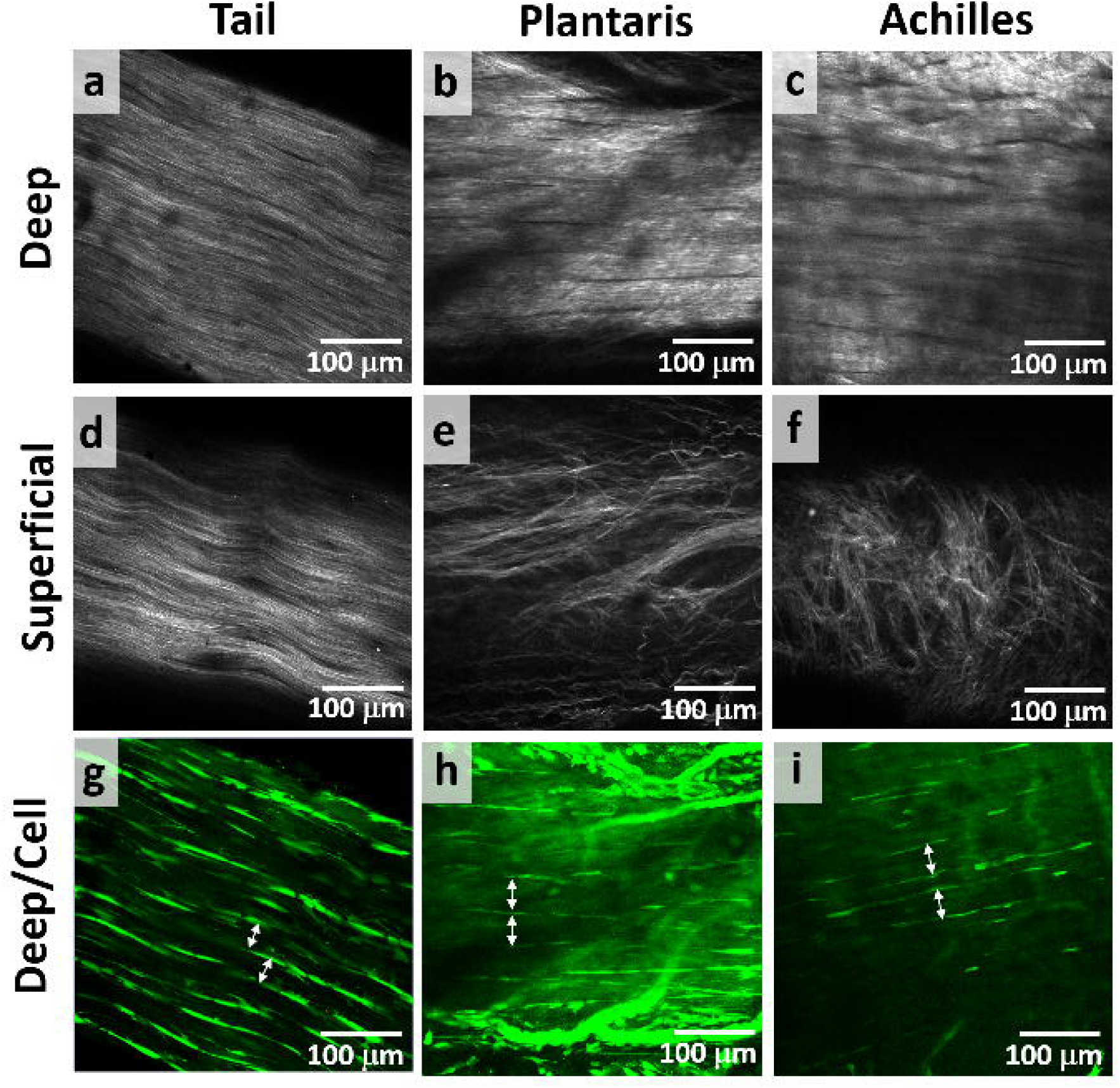

### Three-dimensional (3D) Evaluation of Tendon

The confocal image stacks were reconstructed to view the 3D tendon structure (Fig 5). The 3D reconstruction confirms that cells, shaded in green, surround the collagen bundles (grey) that match the size of the fiber that was identified by histology (Fig 2, 3). For the tail tendon, we again observed no peritenon (Fig 5a), but we observed peritenon for the plantaris and Achilles tendons. The peritenon was populated with blood vessels and localized cells with different morphology than the elongated cells between fibers, shaded in red (Fig 5b, c). In the transverse direction, the 3D structure of tendon shows that cells surround the fiber, highlighted with yellow arrows (Fig 5d-f). Similarly, the cells surround the fiber in an oblique view, highlighted with yellow arrows (Fig 5g-i). Again, we observed no additional boundary. The video of the 3D reconstruction is included as a supplemental video (S.M 1-3). Thus, the 3D rendering of tendon confirms that the definition of a fiber is a bundle of collagen that is outlined by elongated cells, supported by both histologically processed and unprocessed tendons. In all rat tendons studied, the fiber diameters were consistently 10-50 µm.

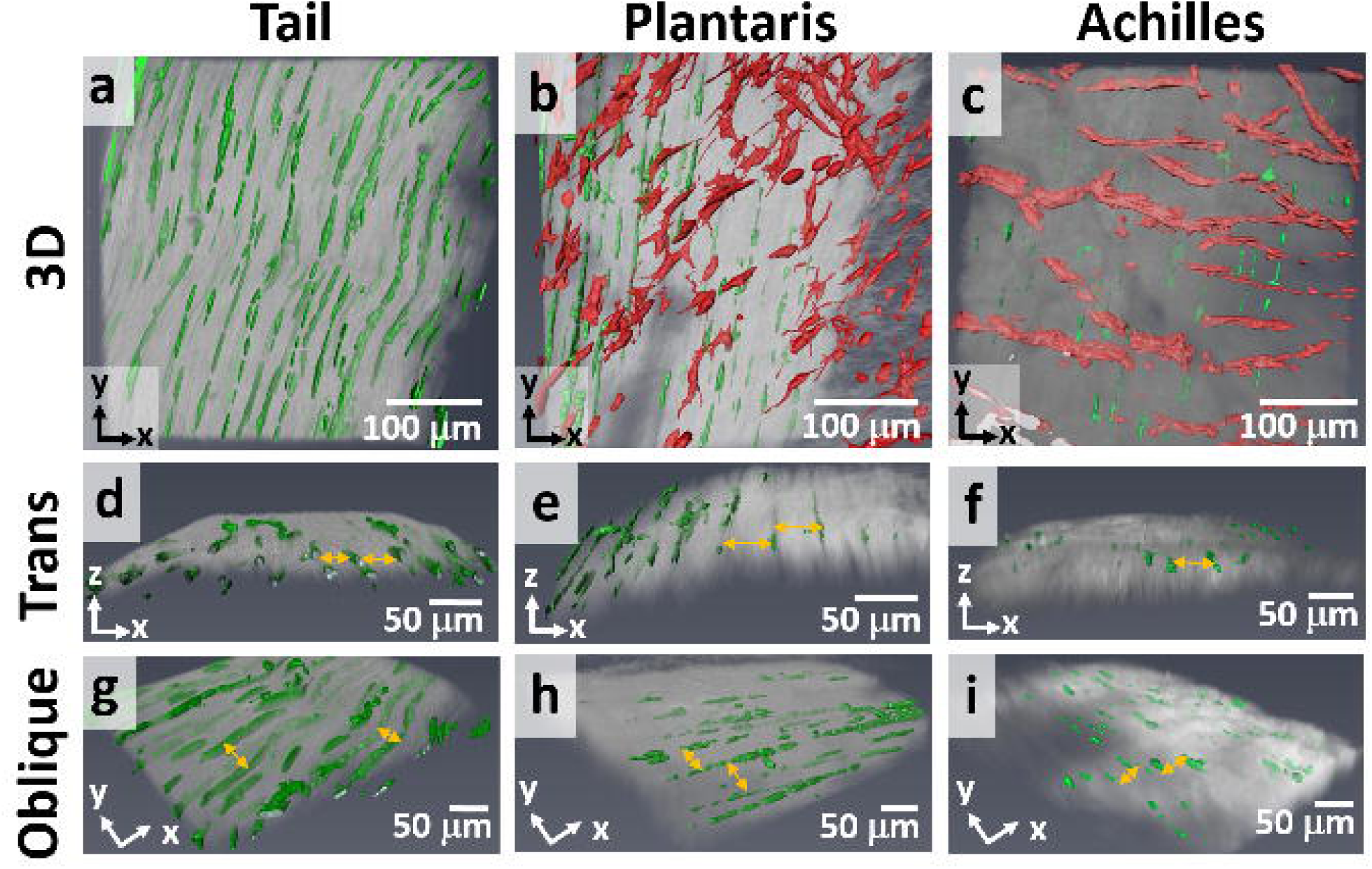

There were no additional boundaries that further divided any of the rat tendons, and we did not observe any evidence of interfascicular matrix, which suggests that there are no fascicles in the rat tendons. A fascicle is defined by a surrounding interfascicular matrix, which is populated with round cells.^17,18^ A fiber may be the largest subunit in rat tendons, and this diameter was consistent in transverse and longitudinal histology and SEM.

## Discussion

In this study, we evaluated the hierarchical structure of rat tendons at multiple length scales for tendons with varying mechanical functions, including the tail, plantaris, and Achilles tendons and created tendon-specific schematics based on our observations (Fig 6). Using multiple imaging methods, we confirmed that rat tendons do not contain fascicles, and thus the fiber is the largest tendon subunit in the rat. In addition, we provided a structurally-based definition of a fiber as a bundle of collagen fibrils that is surrounded by elongated cells, and this definition was supported by both histologically processed and unprocessed tendons. In all rat tendons studied, the fiber diameters were consistently 10-50 µm.

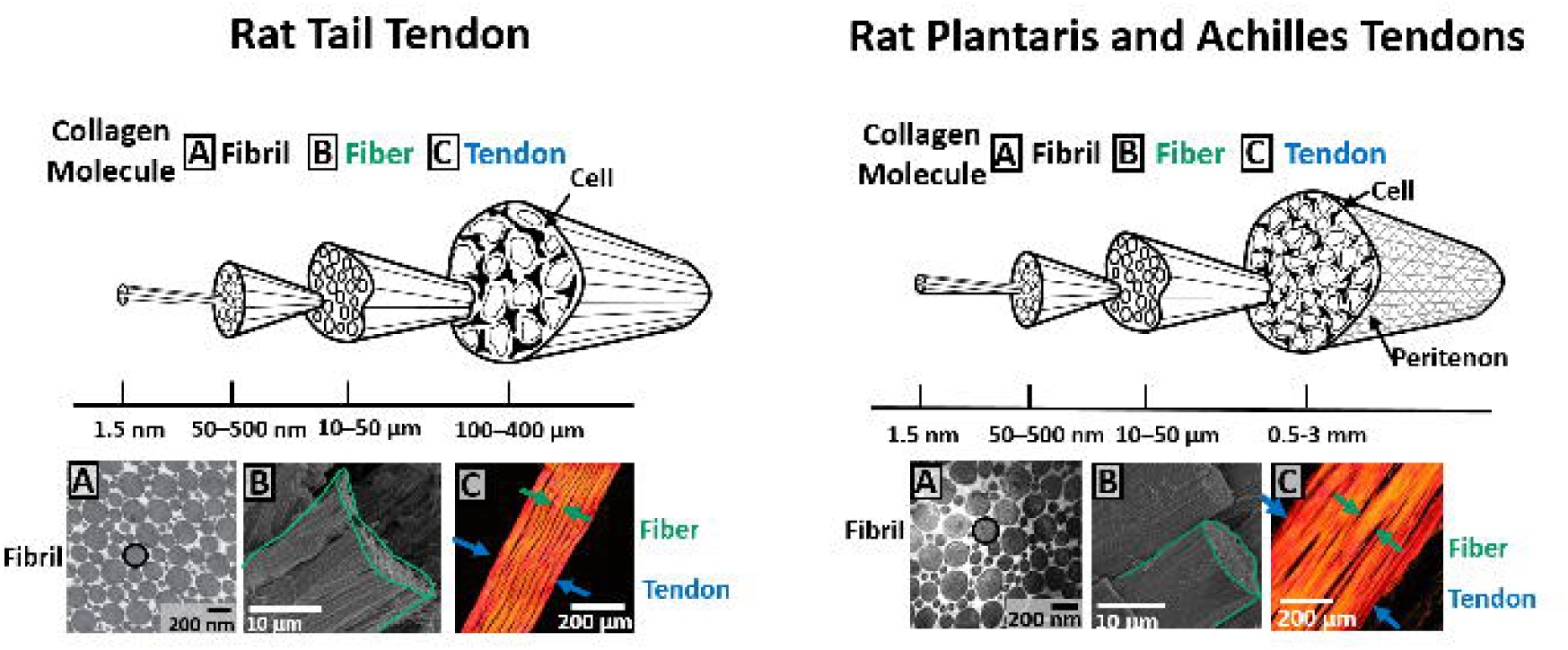

### Structure-based Definitions of the Fascicle and Fiber

This study provides a hierarchical schematics of rat tendons that clarify structural-based definitions of fibers and fascicles (Fig 6). A fascicle is defined by the surrounding interfascicular matrix, which is populated with round cells^17,18^ and is not simply a bundle of fibers. We showed that there are no fascicles in rat tendons, rather the fiber is the largest subunit. Considering the size of rat tendon, it is not surprising that the structural complexity is reduced with the scale of the animal.^15^ In large animals, the fascicle and interfascicular matrix are easily identified and have diameters that are larger than the entire rat tendons.^13–15,20,21^ In the rat tendons studied here, the fiber diameter was 10-50 µm. While this is not the first paper to identify the fiber in rat tendons,^2^ this study provides evidence and a definition of a fiber that is based on characteristic features, confirmed by multiple imaging methods. Importantly, this definition is consistent with the developmental process of tendon. At later stages of development, as cells deposit more collagen, the tendon fiber is delineated as cells are pushed to the periphery.^27^ To avoid confusion, the term fiber should be reserved to specifically identify this hierarchical structure of tendon, not to refer generally to a bundle of collagen.

As a point of clarification, the visualizations of cell density in confocal images and 3D rendering can be misleading because they depend on staining intensity. Because blood vessels and cells on peritenon that can also uptake the fluorophore, less fluorophore penetrated to the deeper planes for plantaris and Achilles tendons. In addition, the diameters of plantaris and Achilles tendons are larger than that of the tail tendon, and thus it contributes to the lack of fluorophore penetration. While it may appear that cell density in the tail tendon is higher than plantaris and Achilles tendons, this is strictly due to the penetration of fluorophore, and this is the cause for lack of cells visualized for Achilles tendon.

### Comparative Anatomy Across Species

We considered if the fiber and fascicle size may be conserved across species. Tendons from larger animals such as horse and human have a subunit that falls within the fiber diameter and is outlined by cells and matches our diameter range, confirming that these tendons have fibers.^17,46,47^ For example, the equine tendon fiber diameter is 1-20 µm^15,29^ and the human tendon primary fiber bundle diameter is 15 µm^21^, which are similar to the fiber diameter 10-50 µm reported here. Thus, the definition of fiber can be applied across species, and this supports the idea that the organization of rat tendon is not a simple scale-down of larger animals, but that the fiber size is *conserved* across species. It is likely that mouse tendon only has fiber, not fascicle, similar to rat tendon due to its small size. Tendon cell volumes^48^ also appear to be consistent across species, which suggests that cells and fibers are the key units in growing aligned collagen region during the developmental process. It is unclear if the fascicle diameters are consistent across species, and it is possible that the number of fascicles or the size of fascicles may scale with the size of the animal.

The absence of fascicles in rat tendons should impact a researcher’s animal model choice: studies that are focused on the structure and mechanics of the interfascicular matrix should exclude rat models. Interestingly, the age-related tendinopathy is mostly attributed to degeneration of the interfascicular matrix^49,50^ and animals that have an interfascicular matrix, such as horses and human, develop naturally occurring age-related tendinopathy^51^, while rats do not appear to do so. While this difference may be due to the animal’s life-span and activity level, studies of age-related tendinopathy should be aware that rats lack fascicles, and thus rats lack interfascicular matrix. Nonetheless, the rat tendon is a useful model system when studying tendon structure and mechanics because rat tendon is a simpler system, especially because the fiber structure and diameter is conserved across species.

### Comparative Anatomy of Rat Tendons

We demonstrated that the anatomy of the rat tail, plantaris, and Achilles tendons have distinct features that are related to their mechanical functions and each tendon can be used to answer specific research questions. The tail tendon’s structure is simple and its mechanics are less variable, making it a suitable model to address fundamental research questions. However, the tail tendon is limited in that it bears low-stresses and lacks the muscle-tendon junction and insertion, cells on peritenon, and blood vessels. Thus, it may not be a suitable model for studying research questions related to high-stress tendons or healing process. The plantaris tendon, on the other hand, bears high-stress in vivo^52^ and has a physiological structure including the muscle-tendon junction and blood vessels. In addition, a synergist ablation, an established rat model applied to study overuse-induced muscle hypertrophy^53–55^, may be a useful model to study tendinopathy. For example, the synergistic ablation model, where the Achilles is removed to overload the synergist plantaris tendon, leads to increased matrix production and thickened plantaris tendon after 4 weeks.^3,4^ This suggests that this model may represent tendinopathic changes. The Achilles tendon also bears high-stress, and we observed that the structure of rat Achilles tendon is a fusion of three tendons, similar to that of human Achilles tendon.^44,45^ Yet, due to its complex structure, the Achilles tendon is difficult to study at multiple length scales. Because of the fusion of three tendons, there is likely inter-tendon sliding^56^ in addition to the micro-scale sliding. This likely contributes to the inhomogeneous loading observed in vivo.^56–58^ This structural complexity should be addressed when using the Achilles tendon as a model system.

While we and others have referred to the rat tail tendon as a ‘fascicle’, this terminology is discouraged. While there is a sheath surrounding rat tail tendons, which one could argue is interfascicular matrix and represents a fascicle. However, this sheath in the tail tendon and the interfascicular matrix of larger animals are distinctly different. The sheath in rat tail tendon can be easily peeled off with no resistance, while the fascicles of larger animals are tightly bound and must be cut to separate.^16^ Thus, rat tail tendons should not be referred to as fascicles and should not be directly compared to fascicles in larger animals.

### Multi-scale Tendon Structure-Function and Damage Mechanics

Tendon structure and mechanics are related at multiple scales, and the fiber scale is likely to be an important contributor to tendon mechanics and damage. For example, tensile damage in the tail tendon is related to the changes in macro-scale mechanical parameters.^32^ Furthermore, in both tail and plantaris tendons, micro-scale sliding, which represents shear between micro-scale structures, is related to both loading^59^ and damage^32^ mechanisms. Because these studies used photobleached lines and optical microscopy, which cannot distinguish between fibrils and fibers, both substructures are captured in the term ‘micro-scale’.^32,59^ Other studies have shown damage that is localized within fibers. Rat tail tendon that is loaded and then labeled with collagen hybridizing peptide (CHP), which binds to denatured collagen, showed a localization of CHP staining in a structure that had the same diameter as fiber observed here.^60^ A similar observation was made in rat flexor carpi ulnaris tendon after mechanical loading, where CHP was localized in a structure with the same diameter as fiber.^61^ In addition, the 3D reconstruction of SHG signals showed that a fiber space between fibers increased in loaded rat patellar tendon, suggesting that damage is related to structural changes in fiber.^37^ Therefore, the fiber is likely to be a key substructure affected during tendon damage.

In summary, this study evaluated the hierarchical structure of rat tail, plantaris, and Achilles tendons at multiple length scales. Structurally-based definitions of fascicles and fibers were established using multiple imaging methods. We confirmed that rat tendons do not contain fascicles and fiber diameters were 10-50 µm in all rat tendons studied. This fiber diameter is conserved across larger species. Specific recommendations were made for the strengths and limitations of each rat tendon as tendon research models

## Acknowledgment

This research was supported and made possible by National Institutes of Health grant No. R01EB002425, National Institute of General Medical Sciences grant No. P20 GM103446, the National Science Foundation grant No. IIA-1301765, National Institutes of Health shared instrumentation grant No. S10 RR027273, and the State of Delaware. We thank the Bioimaging Center at the Delaware Biotechnology Institute and Dr. Jeff Caplan for supporting data acquisition. We thank Francis Karani for providing rats for tendon.

## Authors Contribution

A.H.L. and D.M.E. designed experiments. A.H.L. performed experiments and analyzed data.

A.H.L. and D.M.E. wrote the manuscript.

